# Leukocyte-derived High-mobility group box 1 controls innate immune responses against Listeria monocytogenes

**DOI:** 10.1101/797902

**Authors:** Annika Volmari, Katharina Foelsch, Karsten Yan, Minyue Qi, Karlotta Bartels, Stephanie Kondratowicz, Marius Boettcher, Masahiro Nishibori, Keyue Liu, Robert F. Schwabe, Ansgar W. Lohse, Samuel Huber, Hans-Willi Mittruecker, Peter Huebener

## Abstract

High-mobility group box 1 (HMGB1) is a damage-associated molecular pattern with key proinflammatory functions following tissue injury. Moreover, HMGB1 neutralization was shown to alleviate LPS-induced shock, suggesting a role for the protein as a master therapeutic target for inflammatory and infectious diseases. Here, we report that HMGB1 neutralization impedes immune responses to *Listeria monocytogenes*, a wide-spread bacterium with pathogenic relevance for humans and rodents. Using genetic deletion strategies and neutralizing antibodies, we demonstrate that hepatocyte HMGB1, a major driver of post-necrotic inflammation in the liver, is dispensable for pathogen defense during moderately severe infection with listeria. In contrast, antibody-mediated HMGB1 neutralization and HMGB1 deficiency in myeloid cells effectuate rapid and uncontrolled bacterial dissemination in mice despite preserved basic leukocyte functionality and autophagy induction. During overwhelming infection, hepatocyte injury may contribute to increased HMGB1 serum levels and excessive inflammation in the liver, supporting context-dependent roles for HMGB1 from different cellular compartments during infection. We provide mechanistic evidence that HMGB1 from circulating immune cells contributes to the timely induction of hepatic immune regulatory gene networks, early inflammatory monocyte recruitment to the liver and promotion of neutrophil survival, which are mandatory for pathogen control. In summary, our data establish HMGB1 as a critical co-factor in the immunological clearance of listeria, and argue against HMGB1 neutralization as a universal therapeutic strategy for sepsis.

**Author summary:** High-mobility group box 1 (HMGB1) is an abundantly expressed nucleoprotein with signaling properties following secretion or release into the extracellular space. Given its central immune-regulatory roles during tissue injury and LPS-induced septic shock, interventions aimed at HMGB1 signaling have been advocated as therapeutic options for various disease conditions. Here, we show that antibody-mediated HMGB1 neutralization interferes with immunological defense against *Listeria monocytogenes*, a gram-positive bacterium with high pathogenic relevance for rodents and humans, effectuating uncontrolled bacterial growth and inflammation. Using conditional knockout animals, we demonstrate that while leukocyte functionality is preserved in HMGB1-deficient myeloid cells, HMGB1 released in response to *Listeria* triggers hepatic inflammatory monocyte recruitment and activation of transcriptional immune networks required for the early control of bacterial dissemination. Hepatocyte HMGB1, a key driver of post-necrotic inflammation in the liver, is dispensable for the immune response during moderately severe infection, but likely contributes to excessive hepatitis when infection is uncontrolled and cellular injury is high. We demonstrate a critical and non-redundant role for HMGB1 in the immune-mediated clearance of listeriosis and argue against HMGB1 neutralization as a universal therapeutic option in the context of infection.

## Introduction

Inflammation is an integral component of the host response to infectious and sterile injury of vascularized tissues (1). While the pro-inflammatory functions of distinct molecular signatures of pathogens (pathogen-associated molecular patterns, PAMPs) are well-established and their functions increasingly deciphered, signals that stimulate immune responses under sterile conditions remain enigmatic. It is assumed that molecules released from damaged or injured cells can activate immune effectors, often through shared receptor systems with their PAMP counterparts, to initiate inflammation and wound healing responses (2). Very little is known, however, about the specific contributions and mutual interactions of both classes of molecules in the context of ongoing infection, where pathogen exposure and tissue damage often simultaneously affect immune responses. High-mobility group box 1 (HMGB1) is an abundantly expressed nucleoprotein and considered a prototypical damage-associated molecular pattern (DAMP) with key roles in the initiation of post-necrotic inflammation in various tissues including the skin (3), liver (4,5), pancreas (6,7), skeletal and cardiac muscle (8,9), and central nervous system (10). Moreover, neutralization of extracellular HMGB1 was shown to alleviate LPS-induced septic shock and reduce lethality in polymicrobial abdominal sepsis (11–13), indicating that HMGB1 neutralization may be uniformly beneficial during both sterile and infectious inflammatory processes. Consequently, HMGB1 has repeatedly been suggested as a promising therapeutic target for infectious and non-infectious inflammatory diseases (14–16). Recent studies, however, have linked genetic HMGB1 deletion to defective induction of autophagy, resulting in increased vulnerability of experimental animals to lipopolysaccharide challenge and infection (17,18). Here, we investigated the functions of HMGB1 during systemic listeriosis, a paradigmatic mouse model of gram-positive bacterial infection with high relevance for humans and rodents, and tested its suitability as a therapeutic target during infection. To elucidate its role in the immune response to infection, we determined circulating HMGB1 levels and assessed the induction of immune regulatory networks involved in bacterial clearance and hepatic inflammation in different genetic and antibody-mediated HMGB1 deletion models. We performed extensive *in vitro* studies and bone-marrow transfer experiments to outline the roles of leukocyte subsets during infection. Surprisingly, neutralization of extracellular HMGB1 did not alleviate infection, but effectuated impaired bacterial clearance and exacerbated hepatitis in mice. In contrast to previous studies, we did not observe defective induction of autophagy in HMGB1-deleted tissues which may favor impaired intracellular degradation of listeria. Instead, we demonstrate a critical role for HMGB1 derived from liver-resident and, to a lesser extent, bone-marrow derived immune cells in the early hepatic recruitment of inflammatory monocytes to mount inflammatory gene networks and counteract incipient infection. We further demonstrate that HMGB1 from hepatocytes is dispensable for the immunological control during moderately severe infection, but may contribute to inflammation when overwhelming infection aggravates hepatocyte injury. We identify HMGB1 from different cellular compartments as critical context-dependent triggers of host immune responses to listeria, and conclude that HMGB1 neutralization strategies may not uniformly be beneficial for the host, particularly in the context of bacterial infection.

## Results

### Antibody-mediated HMGB1 neutralization impairs antibacterial defense

In light of the reported beneficial effects of HMGB1-targeted interventions during LPS-induced shock and polymicrobial abdominal sepsis (11,12), we aimed to assess the consequences of antibody-mediated neutralization of extracellular HMGB1 during murine infection with *Listeria monocytogenes*, a paradigmatic mouse model of bloodstream gram-positive bacterial infection (19). Daily administrations of well-established HMGB1-neutralizing antibodies (10,20,21), but not isotype-matched controls, failed to confer protection from systemic listeriosis, but instead impaired bacterial clearance, resulting in significantly higher hepatic bacterial burden 72 hours after infection (**Fig. 1A-B**). FACS analysis demonstrated higher hepatic titers of neutrophils, but not monocytes or dendritic cells, following HMGB1 neutralization (**Fig. 1C**), and histologic examination revealed increased accumulation of bacteria and disturbances of tissue architecture and in the livers and spleens (**Fig. 1D-E**) of anti-HMGB1-treated animals. Concomitantly, hepatic gene expression of inflammatory cytokines *Ccl*2, *Ccl*3/*Mip1α, Cxcl*2, *Tnf*α and *Nos*2, but not *Ifn*γ, was elevated following HMGB1 neutralization (**Fig. 1F**), indicating a profound defect in bacterial clearance, but not in the induction of hepatic inflammation, following neutralization of extracellular HMGB1.

**Fig 1:**
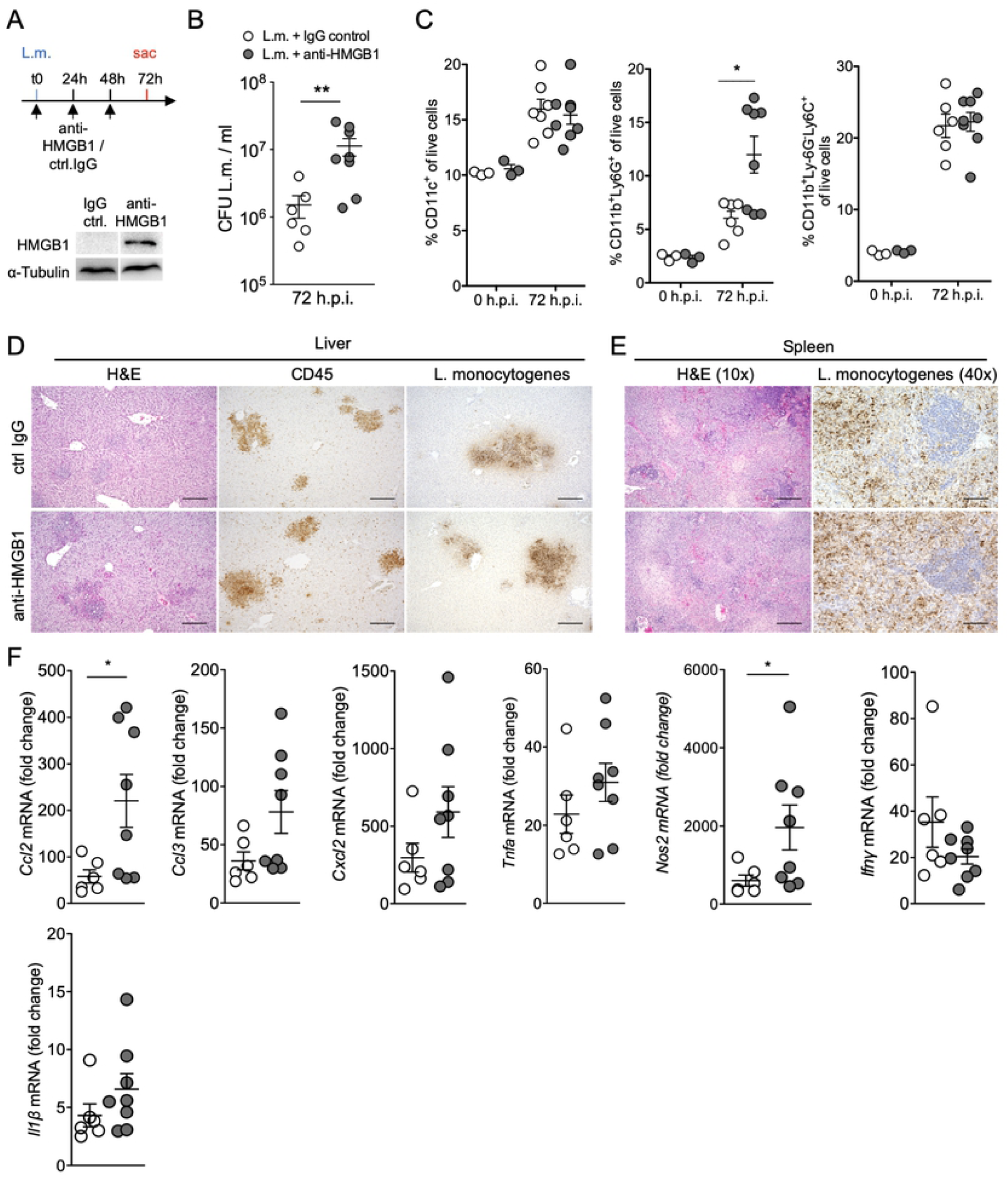
Antibody-mediated HMGB1 neutralization impairs clearance of *Listeria monocytogenes*. (A) Experimental setup and antibody affinity assessment via Western blot analysis of wild-type whole liver lysates. (B) Hepatic titers of *Listeria monocytogenes* in anti-HMGB1-treated and IgG control-treated mice 72 hours after intravenous injection of 2×10^4^ *Listeria monocytogenes* (n=6 and n=8 animals per group, respectively). (C) FACS analysis of CD11c^+^ dendritic cells, CD11b^+^Ly6G^+^ neutrophils and CD11b^+^Ly6C^+^ monocytes 72h after infection. (D) Photomicrographs of hepatic granuloma, infiltrating CD45^+^ cells and *Listeria monocytogenes*. (E) Splenic tissue architecture and accumulation of listeria after 72 hours. (F) qPCR analysis of hepatic expression of key proinflammatory genes 72 hours after infection with listeria. L.m. = *Listeria monocytogenes*. H.p.i. = hours post-infection. Bars = 50 μm. (C): Kruskal-Wallis test with Dunn’s post-test, (F): Mann-Whitney test. * p<0.05, ** p<0.01.

### Hepatocyte HMGB1 is dispensable for antibacterial immunity during moderately severe infection

During systemic listeriosis, bloodborne bacteria disseminate into hepatic and splenic phagocytes and, in the liver, subsequently enter hepatocytes, where they trigger protective immune responses (19). In line with its putative role as a DAMP, we and others have previously demonstrated that hepatocyte HMGB1 not only acts as a key driver of post-necrotic inflammation in the liver (4,5), but also triggers maladaptive wound healing responses during chronic hepatitis (22,23). In contrast to the immuno-stimulatory of hepatocyte HMGB1 in these mostly sterile injury models, hepatocyte-specific HMGB1 deletion via albumin-cre (24) (*Hmgb1*^Δhep^, **Suppl. Fig. S1**) did not affect immune cell recruitment, microabscess and granuloma formation, inflammatory gene induction or bacterial clearance in the first 72 hours following intravenous injection of 2×10^4^ *Listeria monocytogenes* (**Fig. 2A-D**). In fact, despite highly efficient genetic HMGB1 deletion from hepatocytes (**Fig. 2E** and **Suppl. Fig. S1C-E**), circulating HMGB1 levels were low and unaffected by the hepatocyte HMGB1 status (**Fig. 2F**). Immunohistochemistry did not reveal HMGB1 translocation into the cytoplasm of *Hmgb1*^f/f^ hepatocytes (**Fig. 2G**), a feature typically associated with HMGB1 secretion (25). Thus, HMGB1 from injured hepatocytes acts as a key driver of inflammation following sterile liver injury, but is not primarily required to initiate immune responses during moderately severe infection with *Listeria monocytogenes*, when tissue damage is low. While the majority of immune cells constituting hepatic granuloma were strongly HMGB1-positive, a significant fraction displayed reduced or even absent immunoreactivity for HMGB1 (**Fig. 2G**), indicating HMGB1 secretion or passive release, and suggesting a role for myeloid-cell derived HMGB1 in the immune response to listeria.

**Fig 2:**
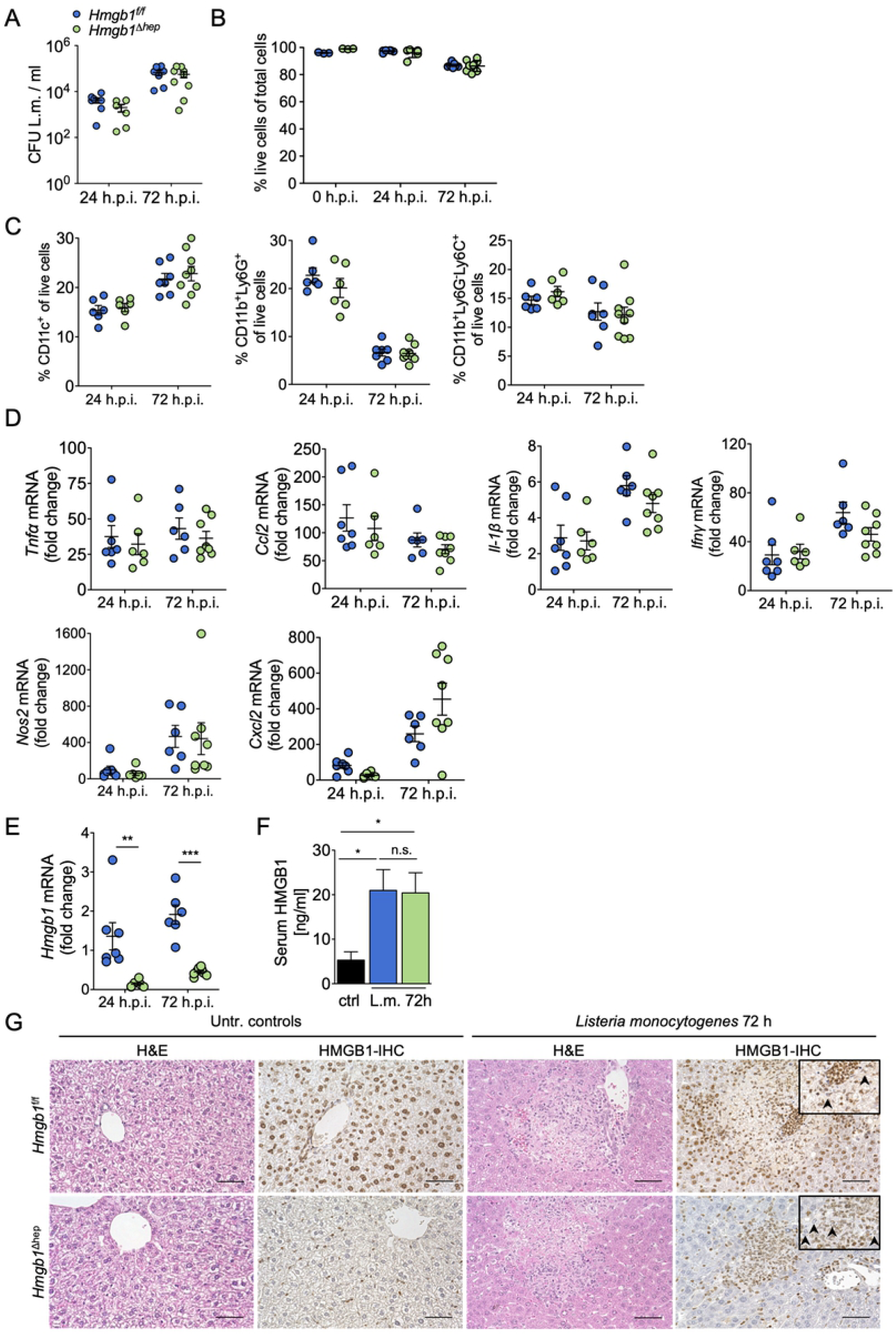
Hepatocyte HMGB1 is dispensable for anti-bacterial immunity. (A) Hepatic titers of *Listeria monocytogenes* in *Hmgb1*^f/f^ and *Hmgb1*^Δhep^ 24 hours and 72 hours after infection (n=6-8 animals per group, respectively). (B) FACS analysis of cellular viability in whole liver cell suspensions. (C) FACS analysis of live hepatic CD11c^+^ dendritic cells, CD11b^+^Ly6G^+^ neutrophils and CD11b^+^Ly6C^+^ monocytes in *Hmgb1*^f/f^ and *Hmgb1*^Δhep^ mice 72 hours after infection. (D) qPCR analysis of hepatic proinflammatory *Tnfα, Ccl2, Il1β, Ifnγ, Nos2* and *Cxcl2* in *Hmgb1*^f/f^ and *Hmgb1*^Δhep^ mice 24 hours and 72 hours after injection of 2×10^4^ *Listeria monocytogenes*. (E) *Hmgb1* mRNA expression in whole liver lysates 24 hours and 72 hours after infection. (F) Serum HMGB1 levels after 72 hours. (G) Microphotographs of liver sections with H&E- and HMGB1-immunostainings, respectively. Arrowheads indicate HMGB1-negative infiltrating immune cells within hepatic granuloma. H.p.i. = hours post-infection. (D), (E): Kruskal-Wallis test with Dunn’s post-test. n.s. = statistically non-significant, *p<0.05, **p<0.01, * * *p<0.001.

### HMGB1 from myeloid cells orchestrates the immunological clearance of Listeria monocytogenes

Kupffer cells are the primary sequestration site of circulating listeria, and phagocytized bacteria were recently shown to induce necroptosis of liver-resident macrophages, triggering type 1 and 2 immune responses that mediate bacterial clearance and the coordinated return to homeostasis (19,26). To test the role of HMGB1 from myeloid cells in the immunological control of systemic listeriosis, we employed myeloid-cell specific HMGB1 ablation via LysM-Cre, effectively deleting HMGB1 from Kupffer cells, monocytes, neutrophils and a minority of dendritic cells, but not B- or T-lymphocytes (*Hmgb1*^ΔLysM^, **Suppl. Fig. S1D-G**) (27). In striking contrast to *Hmgb1*^Δhep^ animals, *Hmgb1*^ΔLysM^ mice displayed profound defects in bacterial clearance beginning as early as 24 hours after infection, ultimately leading to overwhelming infection with accumulation of an ∼100-fold higher bacterial titer in the liver, excessive cell injury, accentuated granuloma formation, and increased hepatic recruitment of myeloid cells, particularly granulocytes, after 3 days (**Fig. 3A-D**). Comparable to anti-HMGB1-treated mice, we also observed increased accumulation of listeria in *Hmgb1*^ΔLysM^ spleens (**Suppl. Fig. S2A-B**) suggesting defective systemic clearance of the pathogen. During overwhelming infection with excessive accumulation of bacteria and exacerbated tissue damage in the liver, hepatocytes from *Hmgb1*^ΔLysM^ animals consistently displayed strong dislocation of nuclear HMGB1 into the cytoplasm (**Fig. 3E**), potentially reflecting increased hepatocyte stress and/or injury. As cytoplasmic HMGB1 translocation typically precedes active secretion, hepatocyte HMGB1 likely contributes to the higher levels of circulating HMGB1 observed in the sera of *Hmgb1*^ΔLysM^ animals after 3 days (**Fig. 3F**) and may contribute to increased inflammation under these circumstances. Transcriptional induction of proinflammatory genes *Tnfα, Nos2, Cxcl2* and *Il1β*, but not *Ifnγ*, was significantly elevated in the livers of *Hmgb1*^ΔLysM^ animals compared to *Hmgb1*^*f/f*^ 72 hours after infection (**Fig. 3G**). Considering only live cells for FACS analysis, we observed comparable hepatic numbers of inflammatory monocytes and dendritic cells, but reduced numbers of neutrophils in *Hmgb1*^ΔLysM^ livers 72 hours after infection (**Fig. 3H**). TUNEL staining confirmed that apoptotic cells were mostly located within and in close proximity to hepatic granuloma of *Hmgb1*^ΔLysM^ animals, whereas very few immune cells were TUNEL-positive in *Hmgb1*^f/f^ controls (**Fig. 3I**). At the same time, listeria were predominantly localized within granuloma, mainly consisting of granulocytes and monocytes, and we did not observe increased numbers of listeria in surrounding HNF4α^+^ hepatocytes of *Hmgb1*^ΔLysM^ mice (**Fig. 3J**). Our findings indicate that in *Hmgb1*^ΔLysM^ animals, increased induction of neutrophil apoptosis or defective clearance of apoptotic granulocyte cell bodies constitutes a characteristic hallmark of the disease phenotype, which may functionally contribute to the defective clearance of listeria (28).

**Fig 3:**
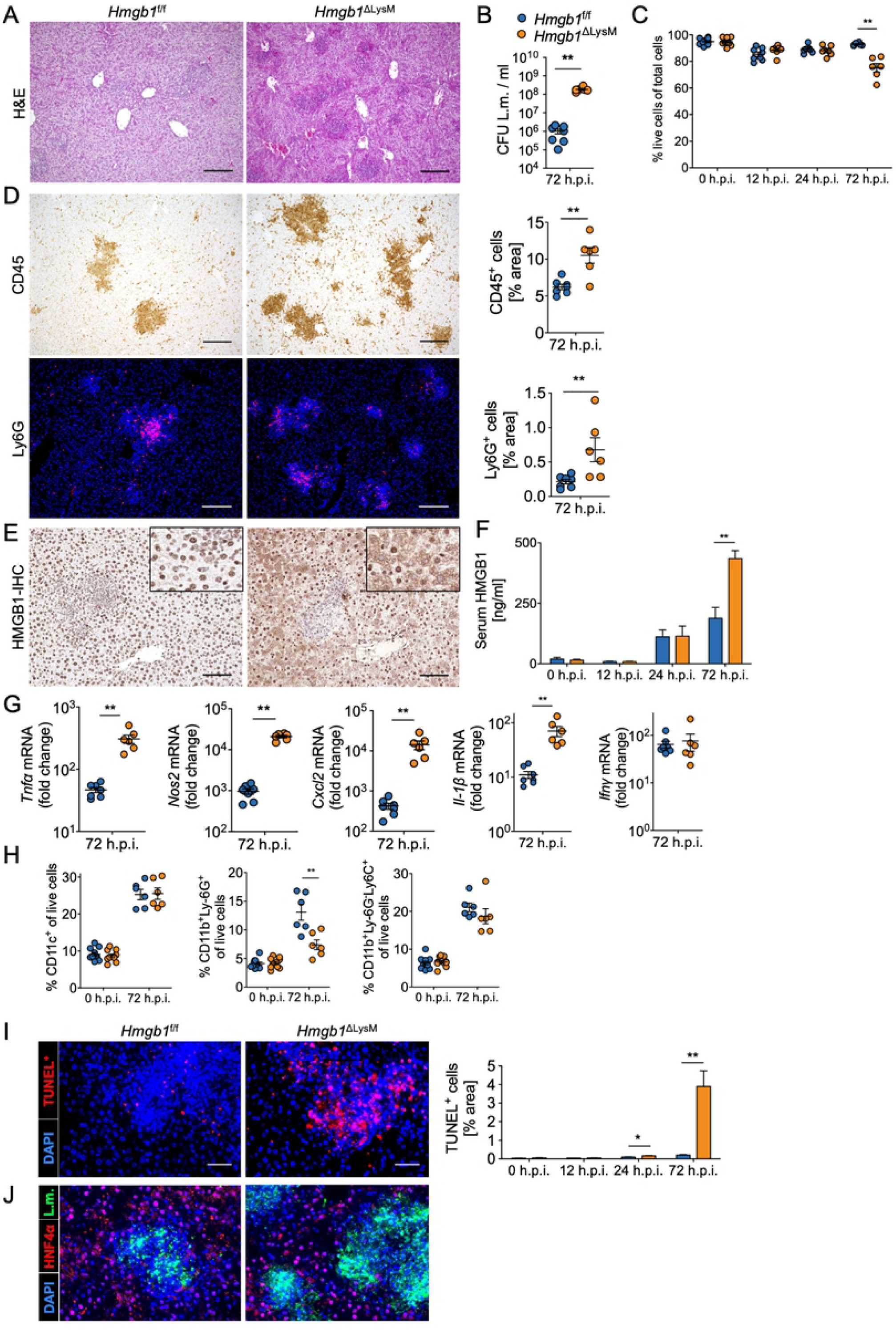
Myeloid-cell derived HMGB1 is required for clearance of listeriosis. (A) H&E stainings of liver sections from *Hmgb1*^f/f^ and *Hmgb1*^ΔLysM^ mice 72 hours after injection of 2×10^4^ *Listeria monocytogenes* (n=6-9 animals per group, respectively). (B) Hepatic bacterial titers in both groups of mice. (C) FACS analysis of cellular viability of whole liver cell suspensions. (D) Immunohistochemistry of hepatic CD45^+^ immune cells and Ly6G^+^ neutrophils. (E) HMGB1 immunostaining of the liver (72 hours) and (F) HMGB1 serum levels assessed by ELISA in *Hmgb1*^f/f^ and *Hmgb1*^ΔLysM^ mice after infection. (G) qPCR analysis of key proinflammatory gene expression in the livers of indicated experimental animals. (H) FACS analysis of live CD11b^+^Ly6G^+^ neutrophils, CD11b^+^Ly6C^+^ monocytes and CD11c^+^ dendritic cells 72 hours after infection. (I) TUNEL staining and J. immunostaining for *Listeria monocytogenes* (green) and HNF-4α (marking hepatocytes, red) in liver sections of the indicated experimental animals. Bars = 50 μm. (B), (D)-(H): Mann-Whitney test. (C): Kruskal-Wallis test with Dunn’s post-test. *p<0.05, * *p<0.01, * * *p<0.001, n.s.=statistically non-significant. H.p.i. = hours post-infection, CFU=colony-forming units.

### Preserved autophagy induction and bactericidal functionality in HMGB1-deficient leukocytes following pathogen exposure

Previous studies have demonstrated associations of HMGB1 and induction of autophagy under conditions of cellular stress (29,17), which were held responsible for various disease phenotypes in HMGB1-deficient experimental animals (30–32). In fact, our observation of increased susceptibility of mice carrying myeloid-cell specific HMGB1 deletion to listeriosis has been linked to impaired autophagy in HMGB1-deficient phagocytes (18). We previously did not detect defective autophagy in a variety of HMGB1-deficient tissues, and did not observe any of the phenotypic features typically associated with impaired autophagy in our experimental mice (33). We thus re-assessed autophagic responses in the present infection model. During infection with listeria, we did not observe differences in the conversion of LC3-I to LC3-II or in the overall induction of LC3-II in the liver over time (**Fig. 4A-D**). Both methods are widely used to assess autophagic flux (34), and were previously applied to link HMGB1 to defective autophagy. We did, however, observe robust accumulation of p62/SQSTM1, an autophagy receptor whose accumulation is linked to impaired autophagic flux, in whole liver lysates of *Hmgb1*^ΔLysM^ particularly after three days of infection. Immunohistochemistry revealed comparable p62 expression in hepatic granuloma of both mice, but strong p62 accumulation in liver parenchyma outside granuloma, indicating that hepatic p62 accumulation is not due to a direct, HMGB1-related autophagy defect in myeloid cells, but likely a reaction of hepatocytes to excessive bacterial burden and inflammation in *Hmgb1*^ΔLysM^ animals. In fact, induction of p62 expression (rather than inhibition of its degradation) has been described in other infectious disease settings (35,36), and may constitute a defense mechanism aiding in the direction of bacteria to autophagic degradation (37) in hepatocytes. In the same line, we observed near identical p62 accumulation in extracts from *Hmgb1*^ΔLysM^ BMDMs and *Hmgb1*^f/f^ BMDMs after infection with *Listeria monocytogenes* (**Fig. 4F-G**) *in vitro*, effectively ruling out cell-intrinsic autophagy defects in *Hmgb1*^ΔLysM^ immune cells. Given the profound impairments of bacterial clearance in *Hmgb1*^ΔLysM^ animals, we next tested bactericidal activities of isolated polymorphonuclear granulocytes (PMNs) as well as primary monocytes, two main effector cell types of innate anti-bacterial immunity (26,38), which are targeted by LysM-Cre (27), from *Hmgb1*^f/f^ and *Hmgb1*^ΔLysM^ mice. We observed a similar ∼40-50% reduction of listeria in the presence of either *Hmgb1*^f/f^ or *Hmgb1*^ΔLysM^ PMNs *in vitro* after 4 hours (**Fig. 5A**), indicating that intracellular HMGB1 is not necessary for PMNs to exert their bactericidal effects after pathogen exposure. Live *Listeria monocytogenes* induced comparable degrees of PMN membrane disintegration and apoptosis, as both numbers of Zombie^+^AnnexinV^-^ PMN (indicating non-apoptotic cell death) and Zombie^+^AnnexinV^+^ (indicating cell death from apoptosis) PMNs tripled after exposure to listeria *in vitro. Hmgb1*^ΔLysM^ PMNs displayed higher numbers of Zombie^-^ AnnexinV^+^ (indicating early apoptosis) PMNs after isolation, suggesting that while listeria regularly induces apoptotic and non-apoptotic cell death in these cell types, HMGB1 may prevent early programmed cell death events in PMNs (**Fig. 4B**). Apart from the overwhelming bacterial burden *in vivo* at later time points in *Hmgb1*^ΔLysM^ animals, it is thus conceivable that HMGB1 may act as a paracrine survival signal, effectuating the excessive accumulation of apoptotic PMNs observed in the absence of leukocyte HMGB1.

**Fig 4.**
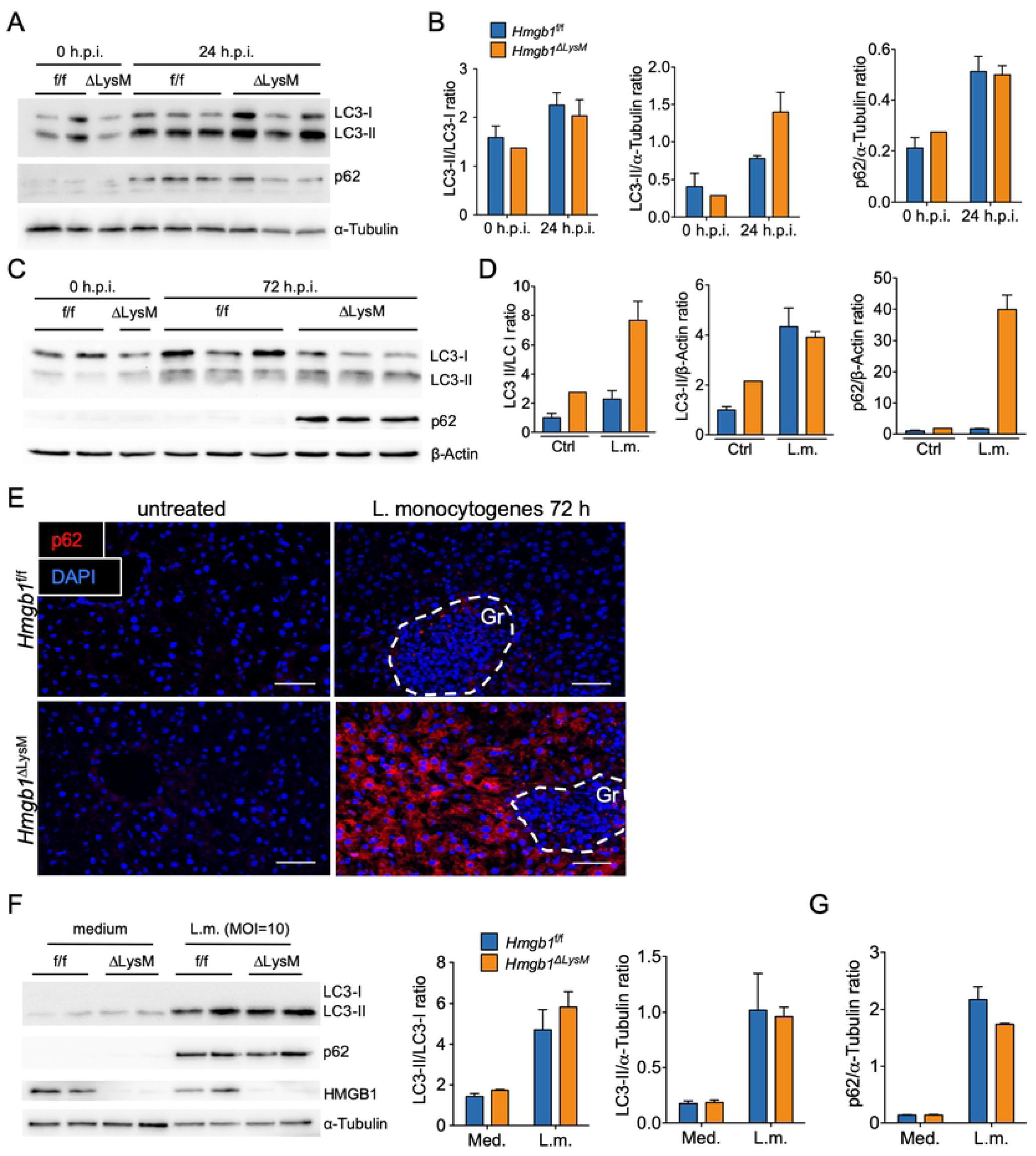
Autophagy induction in leukocytes occurs independent of HMGB1 status. (A) Western blotting of LC3-I, LC3-II, p62 and α-tubulin expression in whole liver extracts of *Hmgb1*^f/f^ and *Hmgb1*^ΔLysM^ mice at baseline and 24 hours after infection with 2×10^4^ *Listeria monocytogenes*. (B) Densitometry analysis. (C) Western blotting of aforementioned proteins after 72 hours and (D) corresponding densitometry analysis. (E) Immunofluorescence staining for p62 expression on liver cryosections 72 hours after infection. (F) Western blot analysis and (G) Corresponding densitometry of LC3-I, LC3-II, p62, HMGB1 and α-tubulin expression of primary isolated BMDMs after *in vitro*-infection with live *Listeria monocytogenes*. Results are representative of at least three independent experiments. Bars = 50 μm.

**Fig 5:**
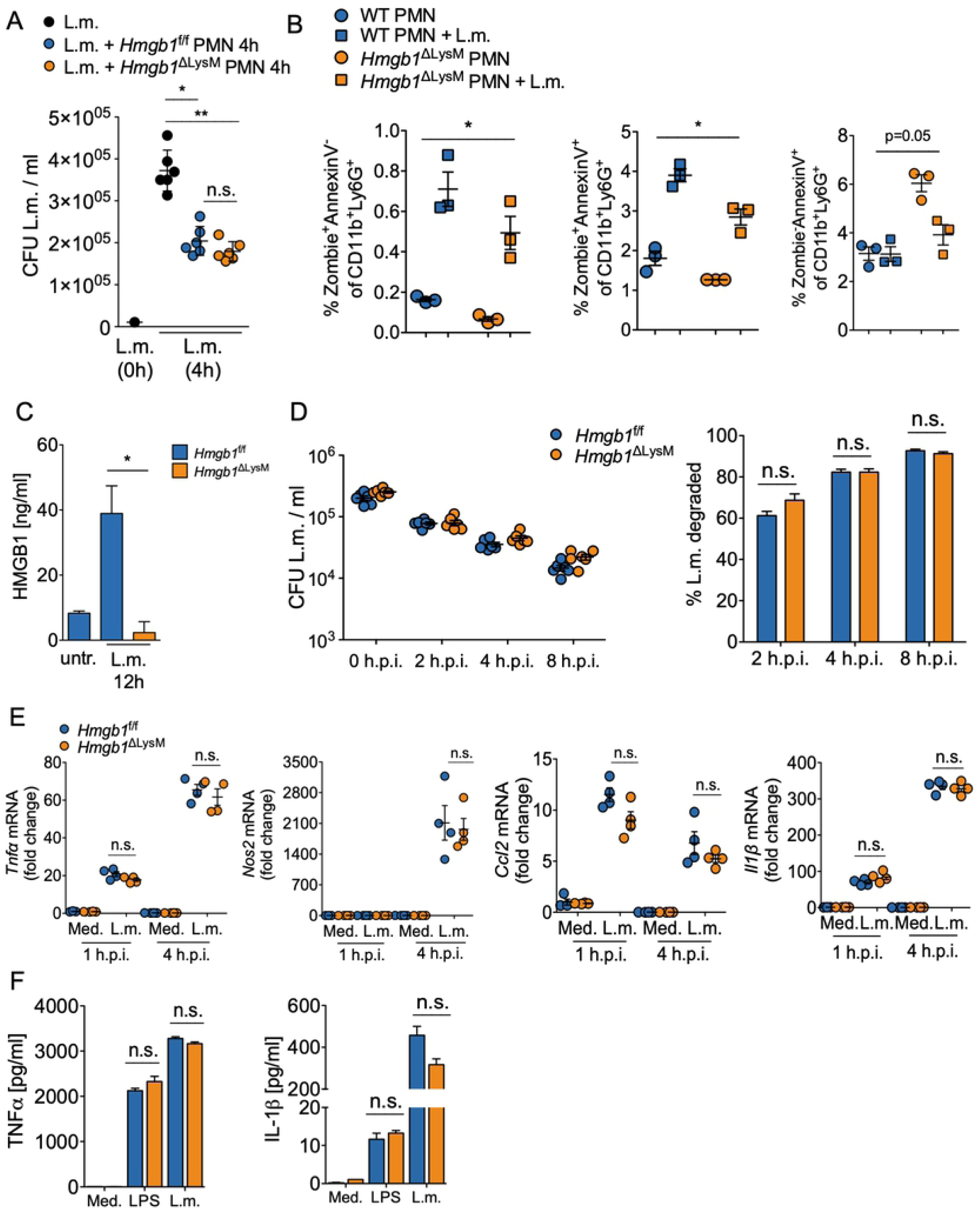
Preserved leukocyte bactericidal activity in phagocytes of *Hmgb1*^ΔLysM^ mice. (A) *In vitro*-bactericidal activity of neutrophils from *Hmgb1*^f/f^ and *Hmgb1*^ΔLysM^ mice (MOI=0.05, n=6 setups per group, results representative from at least three independent experiments). (B) FACS-based assessment of neutrophil expression of Zombie and Annexin V for early and late apoptosis *in vitro* after isolation and exposure to *Listeria monocytogenes* (MOI=0.05). (C) HMGB1 levels in supernatants of untreated and listeria-stimulated isolated primary bone-marrow derived macrophages (MOI=10). (D) *In vitro*-assessment of BMDM phagocytosis and intracellular degradation of *Listeria monocytogenes*. (E) qPCR analysis of proinflammatory gene expression of primary BMDMs under the indicated conditions and (F) ELISA for inflammatory cytokines after 4 hours of incubation with LPS (1ug/ml) or *L*.*m*. (MOI=10), respectively. (A), (C), (E): Kruskal-Wallis test with Dunn’s post-test. *p<0.05, * *p<0.01, n.s.=statistically non-significant.

In light of the critical role of hepatic macrophages (both Kupffer cells as well as circulating bone-marrow derived monocytes) as primary sequestration sites of circulating listeria, we next assessed monocyte responses after *in vitro*-exposure to *Listeria monocytogenes*. We observed a near-complete absence of intra- and extracellular HMGB1 in cultivated primary *Hmgb1*^ΔLysM^ macrophages (**Fig. 5C and Suppl. Fig. S1G**). Pathogen uptake into *Hmgb1*^ΔLysM^ BMDMs was comparable to *Hmgb1*^f/f^ BMDMs, and preceded intracellular bacterial degradation independent of the mouse genotype, resulting in >90% degraded listeria eight hours after internalization of bacteria in both groups (**Fig. 5D**). HMGB1 deletion neither affected inflammatory gene transcription in BMDMs nor TNFα or IL1β release at baseline or after exposure to comparable bacterial concentrations (MOI=10) or LPS (**Fig. 5E-F**), thus ruling out cell-intrinsic interferences between our deletion strategy and inflammatory gene induction or cytokine release in BMDMs.

### *Differential mononuclear cell recruitment and immune pathway activation contribute to impaired bacterial clearance in Hmgb1*^ΔLysM^

Having established the apparently preserved basal functions of immune cells from *Hmgb1*^f/f^ and *Hmgb1*^ΔLysM^ mice *in vitro*, we aimed to further elucidate mechanisms that may account for the impaired immunologic control of systemic listeriosis in *Hmgb1*^ΔLysM^ animals. We thus examined immune responses in the early course of infection, where bacterial titers started to diverge between *Hmgb1*^f/f^ and *Hmgb1*^ΔLysM^ animals (**Fig. 6A**). We observed robust early infiltration of neutrophils into the liver, with higher PNM numbers in *Hmgb1*^ΔLysM^ mice after 24 hours, reflecting intact recruitment in both groups, proportional to the respective pathogen burden (**Fig. 6B**). In contrast, we observed a profound reduction of infiltrating CD11b^+^ Ly6G^-^ Ly6C^+^ cells, and particularly CD11b^+^Ly6G^-^Ly6Chigh cells into *Hmgb1*^ΔLysM^ livers (cf. **Suppl. Fig. S8** for gating strategies) contrasting the increased bacterial burden (**Fig. 6C**). In the context of bacterial infection, these inflammatory monocytes are rapidly mobilized from the bone marrow and recruited to the liver and spleen, where they exert important functions in the orchestration of the ensuing immune response (39,40). Moreover, we and others (26) observed a strong accumulation of F4/80-positive cells in the liver 24 hours after infection, which largely disappeared after 3 days as assessed by F4/80 immunohistochemistry and F4/80 qPCR (**Suppl. Figure S3**). The effect is typically attributed to phagocyte necroptosis (26), and, based on hepatic F4/80 expression, was markedly enhanced in *Hmgb1*^ΔLysM^ animals, potentially affecting inter-phagocyte crosstalk.

**Fig 6.**
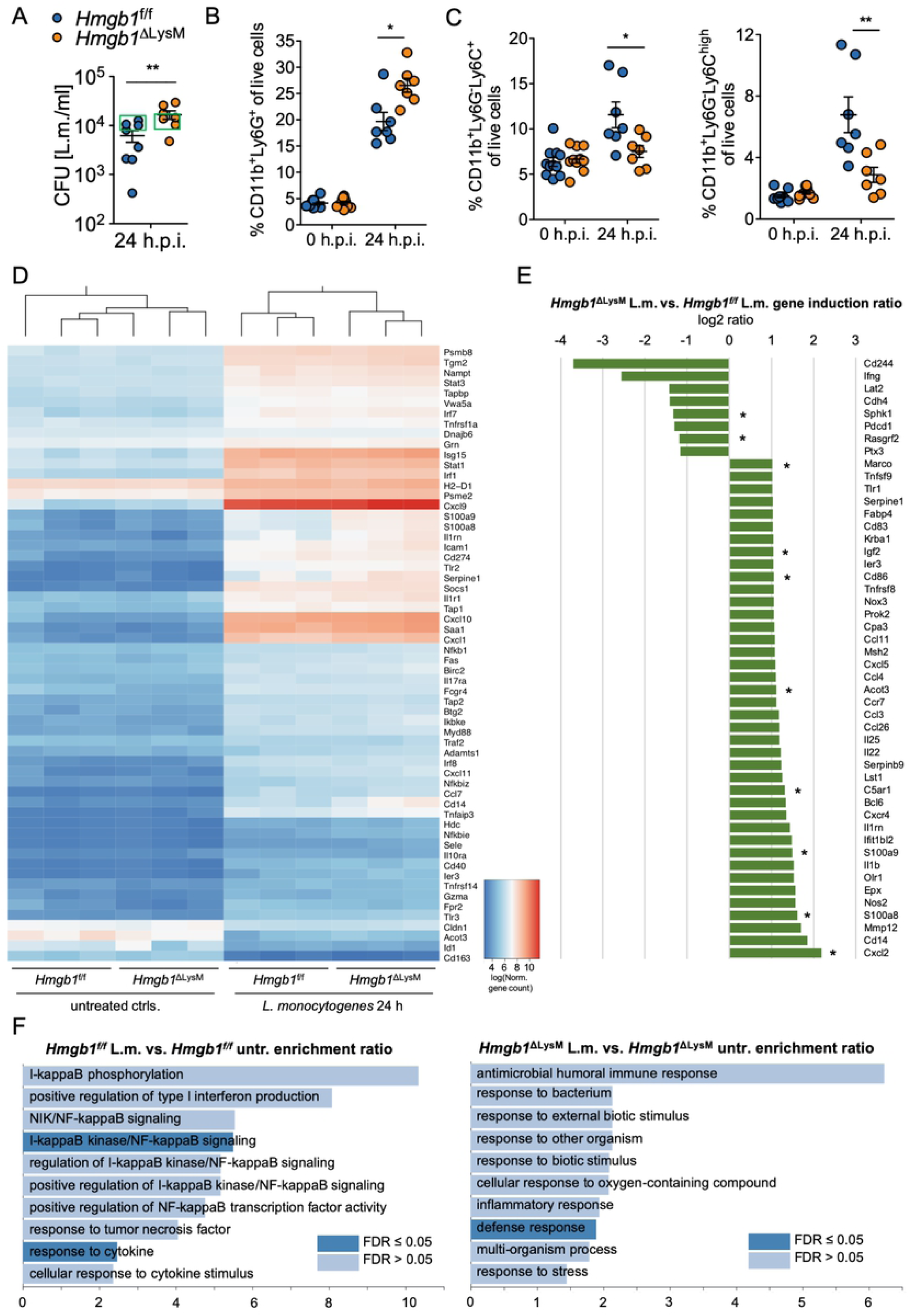
Impaired monocyte recruitment and differential immune pathway activation in *Hmgb1*^ΔLysM^ mice early after infection with *Listeria monocytogenes*. (A) Hepatic bacterial burden in *Hmgb1*^f/f^ and *Hmgb1*^ΔLysM^ mice 24 hours after infection (n=7-8 animals per group). Green boxes indicate mice used for Nanostring analysis. (B) FACS analysis of hepatic CD11b^+^Ly6G^+^ neutrophils and (C) CD11b^+^Ly6C^+^ and CD11b^+^Ly6C^high^ monocytes 24 hours after infection. (D) Heatmap analysis of hepatic gene expression of mice marked in (A), with comparable pathogen burden after 24 hours. (E) List of all tested genes with >1 log_2_ differential expression between infected *Hmgb1*^f/f^ and *Hmgb1*^ΔLysM^ mice. (F) Overrepresentation enrichment analysis of gene induction in *Hmgb1*^f/f^ and *Hmgb1*^ΔLysM^, respectively, after 24 hours of infection. H.p.i. = hours post-infection. FDR = false discovery rate. (A), (E): Mann-Whitney test. (B), (C): Kruskal-Wallis test with Dunn’s post-test. *p<0.05, * *p<0.01.

Given the “receptor promiscuity” of HMGB1, which reportedly associates with partner molecules such as LPS, ssDNA, IL1β and nucleosomes to enhance activation of their respective receptors (41), we next studied innate immune activation patterns in livers of *Hmgb1*^f/f^ and *Hmgb1*^ΔLysM^ animals with comparable bacterial titers and thus equivalent exposure to pathogen and associated PAMPs after 24 hours (**Fig. 6A**, marked green). Heat map analysis demonstrated close transcriptional resemblance between untreated *Hmgb1*^f/f^ and *Hmgb1*^ΔLysM^ mice, whereas treated mice from both groups clearly clustered according to their genotype, indicating differential gene expression depending on the leukocyte HMGB1 status following infection (**Fig. 6D**). Of note, Nanostring analysis revealed >log2-fold differential expression of only 48/734 (6.5%) genes involved in innate immune cell regulation between *Hmgb1*^f/f^ and *Hmgb1*^ΔLysM^ livers, and there was a dominance of proinflammatory gene overexpression in HMGB1-deficient animals (**Fig. 6E**). Overrepresentation enrichment analysis revealed entirely different patterns of immune pathway activation in *Hmgb1*^f/f^ and *Hmgb1*^ΔLysM^ mice despite identical bacterial burden in both groups of mice (**Fig. 6F and Supplemental Figure S6**). *Hmgb1*^f/f^ predominantly displayed induction of I-kappaB kinase/NF-kappaB signaling, which constitute downstream targets of several cell surface receptors and critically mediate DNA transcription, cytokine induction and cellular survival in the context of infection and cellular stress (42). By contrast, *Hmgb1*^ΔLysM^ instead revealed activation of pathways related to pathogen response and induction of humoral antimicrobial responses, and displayed no overrepresentation of IkB/NFkB-related pathways compared to untreated *Hmgb1*^ΔLysM^ controls. Our results indicate that in wildtype animals, HMGB1 may act as a critical endogenous co-activator of receptor-mediated IkB/NFkB activation and subsequent downstream control of the immune response to *Listeria monocytogenes*.

Serum levels of IL1β and IFNγ were comparable between *Hmgb1*^f/f^ and *Hmgb1*^ΔLysM^ 24 hours after infection despite higher bacterial titers in *Hmgb1*^ΔLysM^ at this stage of infection (**Supplemental figure S2C**), raising the possibility that a decrease of distinct soluble mediators of inflammation may account for an inadequate early immune response. In fact, transcriptional induction of *interferon-γ*, a critical mediator of anti-listerial immunity (43,44), was markedly reduced in *Hmgb1*^ΔLysM^ in the Nanostring analysis (**Fig. 6E**) and displayed induction levels comparable to *Hmgb1*^f/f^ after 72 hours of infection, despite >100fold higher bacterial titers in the former group, suggesting a relative shortage of IFNy in *Hmgb1*^ΔLysM^. Of note, when we infected mice with a less virulent batch of bacteria causing attenuated infection with lower bacterial titers, disease phenotypes were comparable in *Hmgb1*^f/f^ and *Hmgb1*^ΔLysM^ mice, with similar hepatic bacterial titers after 72 hours (**Suppl. Fig. S4A**), suggesting that the stress response triggered by HMGB1 is critically important predominantly during severe infection with accentuated cell death. In this model, serum IFNγ levels displayed a moderate reduction in *Hmgb1*^ΔLysM^ (**Suppl. figure S4B**) mice despite comparable bacterial burdens, and reduced infiltration of immune cells into the liver (**Suppl. Fig. S4C-D**). In contrast, qPCR analysis from severely infected livers revealed timely induction of key proinflammatory genes *Tnfα, Nos2* and *Cxcl2*, which later on paralleled the excessive bacterial burden in the hepatic microenvironment, showing that inflammatory gene induction *per se* is not impaired in the absence of leukocyte HMGB1, and later on reflects the immune response to overwhelming infection and/or increased extracellular HMGB1 concentrations (**Fig. 3F**).

### HMGB1 from circulating and liver-resident immune cells contributes to microbicidal activity

Infection with listeria triggers liver-resident phagocyte necroptosis followed by monocyte recruitment and the induction of an antibacterial type 1 inflammatory response (25). To further elucidate the relative contributions of HMGB1 from liver-resident phagocytes and bone-marrow derived immune cells, we generated *Hmgb1*^ΔLysM^ bone-marrow chimeric mice. Both wild-type (WT) mice replenished with *Hmgb1*^ΔLysM^ bone marrow and *Hmgb1*^ΔLysM^ mice with WT bone marrow exhibited impaired clearance of listeria, with an attenuated disease phenotype compared to *Hmgb1*^ΔLysM^>*Hmgb1*^ΔLysM^ chimera (**Fig. 7A-B**). We observed higher bacterial titers, exacerbated hepatic and splenic inflammation and increased expression of hepatic proinflammatory genes in both *Hmgb1*^f/f^ >*Hmgb1*^ΔLysM^ and *Hmgb1*^ΔLysM^>*Hmgb1*^f/f^ after 3 days (**Fig. 7C-D**). Defects in bacterial clearance were more pronounced in WT mice reconstituted with *Hmgb1*^ΔLysM^ bone marrow, suggesting that HMGB1 signaling from circulating myeloid cells towards tissue-resident immune cells is more critical for bacterial defense than vice versa.

**Fig 7:**
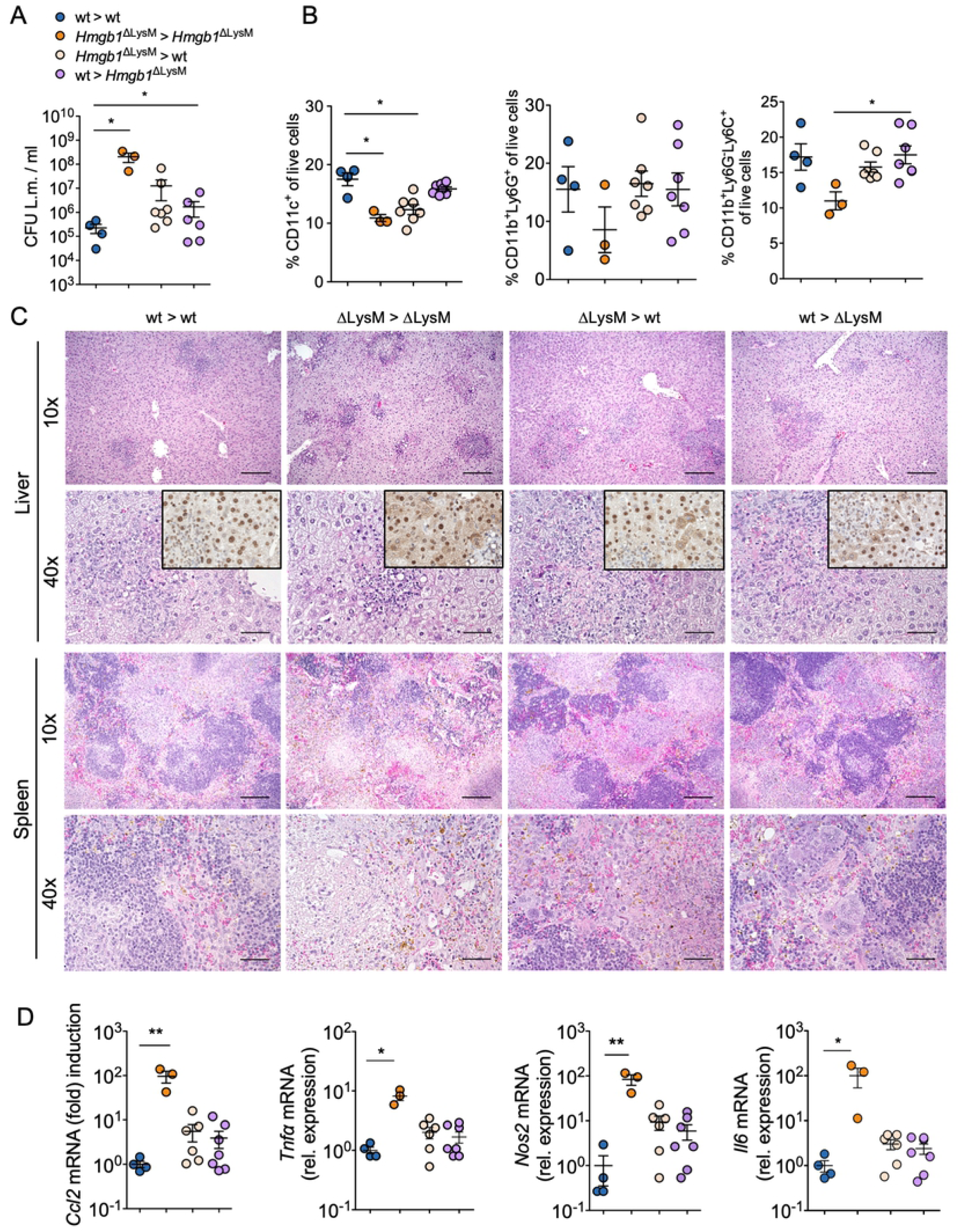
Bone-marrow chimera reveal predominant contribution of HMGB1 from bone-marrow and liver-resident immune cells to successful bacterial clearance. (A) Hepatic bacterial titers of the indicated animals 72 hours after injection of *Listeria monocytogenes* (n=3-7 animals per group, respectively). (B) FACS analysis of hepatic CD11c^+^ dendritic cells, CD11b^+^Ly6G^+^ neutrophils and CD11b^+^Ly6C^+^ monocytes 72 hours after infection. (C) Representative photomicrographs of H&E-staining and HMGB1 immunohistochemistry (inserts) of liver sections of the indicated animals after 72 hours. (D) qPCR of hepatic gene expression of proinflammatory *Ccl2, Tnfα, Nos2* and *Il-6*. (A), (B), (D): Kruskal-Wallis test with Dunn’s post-test. *p<0.05, * *p<0.01. Bars = 200μm.

## Discussion

Inflammation is a critical element of the host response to injury and infection. Whereas molecular signatures of “foreignness” stimulate immune effectors upon pathogen exposure, DAMPs are host molecules released from stressed or decaying cells and initiate inflammation and wound healing responses following tissue damage (2). Intuitively and demonstrably, elevated circulating DAMP levels can also be detected during infections with protozoans (45,46), fungi (47), viruses (48,49) and bacteria (50,51), presumably reflecting context-dependent degrees of tissue damage and DAMP secretion by activated immune and parenchymal cells. The extent of evolutionary conservation and the remarkably high expression levels of the prototypical DAMP candidate HMGB1 in virtually all mammalian tissues (52) suggests important functions for the molecule in health and disease, and previous studies have demonstrated that HMGB1 not only triggers post-necrotic inflammation, but also mediates lethality in LPS-induced septic shock in mice (11,13). In light of encouraging results from animal studies using molecular inhibitors or upstream inhibition of HMGB1 release even in late phases of septic shock, the molecule has gained significant attention as a potential target for therapeutic interventions in affected patients (53,14). In part due to the early postnatal lethality of global HMGB1 knockout animals (54), however, our current knowledge about the functions of HMGB1 *in vivo* remains limited and requires significant advances before translation into clinical medicine. In fact, first reports from mice with conditional genetic HMGB1 deletion revealed detrimental effects during LPS-induced shock and bacterial sepsis (18).

Here, we aimed to address the therapeutic suitability of HMGB1 neutralization during infection with *Listeria monocytogenes*, a widely distributed bacterium causing severe infections predominantly in pregnant women, elderly and immunocompromised patients. Surprisingly, neutralizing HMGB1 antibodies did not confer host protection, but instead resulted in higher bacterial titers, aggravated hepatic inflammation and exacerbated tissue damage, indicating functions of extracellular HMGB1 that are critical for pathogen defense and limit its suitability as a therapeutic target for infections (14–16). The reasons for these divergent findings remain elusive but may be attributed to non-specific or off-target effects of HMGB1 antibody administration, highly context-specific functions of the molecule in different disease scenarios (i.e., gram-negative or gram-positive infection), or even subtle differences in mouse strain genetics or other confounders (i.e., the intestinal microbiome), which notoriously affect experimental outcomes. Of note, however, our findings from antibody-mediated HMGB1 neutralization were confirmed in genetic models of HMGB1 deficiency as demonstrated by a comparable phenotype evoked by myeloid cell-specific HMGB1 deletion during systemic listeriosis. While we cannot completely rule out that the phenotypes of both experimental approaches are mechanistically unrelated, our data suggest that HMGB1-mediated signaling from leukocytes is needed to contain bacterial growth in the early phase of infection. In this concept, genetic or early pharmacologic inhibition of HMGB1 signaling favors overwhelming infection and effectuates PAMP-driven excessive inflammation. The fact that HMGB1 from myeloid cells - but not from hepatocytes - critically regulates the induction of antibacterial immunity during moderately severe infection suggests that professional phagocytes, the primary sites of listeria sequestration in the liver and the spleen (19), release HMGB1 compatible with a “DAMP” role of myeloid cell HMGB1. We demonstrate, however, that the DAMP function of HMGB1 are not restricted to a general promotion of inflammation – in fact, our analysis of inflammatory gene expression in *Hmgb1*^ΔLysM^ and *Hmgb1*^f/f^ livers with the same pathogen burden revealed more up-than downregulated proinflammatory mediators in the absence of leukocyte HMGB1 after 24 hours. Strikingly, our analyses further show that during overwhelming infection, when cell death is more prevalent, HMGB1 translocates into the cytoplasm of injured hepatocytes, likely preceding active secretion (25) and contributing to elevated HMGB1 serum levels. Our findings suggest that HMGB1 from hepatocytes is dispensable when cellular injury is negligible and the initiation of innate immunity is mediated by leukocyte HMGB1. During overwhelming infection, however, hepatocellular injury aggravates, and release of HMGB1 may contribute to excessive inflammation in severely septic animals. These observations are compatible with the observations that hepatocyte HMGB1 triggers damage-induced inflammation in the liver (5), and mediates lethality during severe bacterial sepsis (13). Further studies exploring interventions targeted against specific HMGB1 isoforms, during different phases of the immune response and in varying disease severities may decisively augment our understanding of the dynamics of HMGB1 signaling during infection. In this regard, the fact that ablation of HMGB1 from dendritic cells via Cd11c-Cre (*Hmgb1*^ΔDC^) also increases bacterial titers and inflammation in the liver (**Suppl. Fig. S5**) further corroborates the importance of intercellular signaling via HMGB1 in the induction of an effective antibacterial response.

The general phenotype observed in our *Hmgb1*^ΔLysM^ animals during infection is in accordance with a previous report (18), in which the increased sensitivity of mice with myeloid-cell specific HMGB1 deletion towards LPS challenge and infection with listeria was ascribed to impaired autophagy, reflected by the reduced accumulation of autophagy marker LC3-II, in phagoyctes. Using our conditional HMGB1 knockout animals, we previously found no phenotypic or metabolic alterations typically associated with defective autophagy, in HMGB1-deficient cells including hepatocytes, macrophages, mouse embryonic fibroblasts and cardiac muscle cells at baseline or under conditions of cellular stress (33). Likewise, in the present study, we did not observe autophagy defects in phagocytes of *Hmgb1*^ΔLysM^ mice as a potential explanation for impaired pathogen clearance. We detected comparable accumulation of LC3-II, a marker of autophagy induction, in stimulated primary BMDMs as well as in whole liver lysates. While we did observe accumulation of p62 in whole livers of *Hmgb1*^ΔLysM^ mice, immunohistochemistry revealed high p62 expression almost exclusively in hepatocytes surrounding granuloma, likely as a “stress response” in *Hmgb1*^ΔLysM^ livers exposed to overwhelming bacterial burden. On a more functional level, we observed intact phagocytic activity, intracellular bacterial degradation and inflammatory cytokine induction in HMGB1-deficient monocytes, as well as preserved bactericidal activity of neutrophils in the absence of intracellular HMGB1, indicating that these critical immunological functions are likely induced by PAMPs and/or other DAMPs during listeriosis. However, *Hmgb1*^ΔLysM^ mice exhibited impaired early recruitment of inflammatory monocytes to the liver, a cell population which is mobilized from the bone marrow in the context of bacterial infection in a CCR2-dependent manner (55), and subsequently enters the liver and spleen to orchestrate vital systemic and local innate immune responses (39,40). CCR2-knockout animals display a comparable disease phenotype to *Hmgb1*^ΔLysM^ during systemic listeriosis (55). Thus, impaired induction of this cellular compartment in the early course of infection may account for a crucial immunological disadvantage of *Hmgb1*^ΔLysM^ mice, effectuating disseminating hepatic granuloma formation, accumulation of large numbers of apoptotic polymorphonuclear phagocytes, and ultimately failure to eliminate listeria from the liver. While supporting the notion that HMGB1 released from injured or infected cells may guide specific immune cells to sites of ongoing tissue damage, we also found *in vivo*-evidence for an involvement of HMGB1 as a co-activator of NFkB-dependent signaling, a key pathway of leukocyte reprogramming and immune activation, in the context of systemic listeriosis. In our Nanostring analysis of mice from both groups with equivalent pathogen burden, we observed a robust over-representation of NFkB-related pathways in *Hmgb1*^f/f^ livers after 24h, which was virtually absent in *Hmgb1*^ΔLysM^ animals. Interestingly, mice deficient for MyD88, a downstream adaptor protein of IL1-R and most Toll-like receptors, are similarly susceptible to overwhelming listeriosis, indicating a possible role for HMGB1 as a co-ligand for receptor-mediated MyD88 activation (56). Previous studies, showing that mice deficient for TLR2 or its coreceptor CD14, but not TLR4 or RAGE, are comparably prone to overwhelming listeriosis (56–59) suggests a role for TLR2 in HMGB1-mediated induction of NFkB-signaling. Indeed, several studies have indicated functional HMGB1-TLR2 interactions (60,61). It is further conceivable that extracellular HMGB1 released from myeloid cells enhances neutrophil antibacterial activity, i.e. via induction of neutrophil extracellular trap (NET) release (62). Moreover, HMGB1 may sustain neutrophil survival or partake in the removal of apoptotic neutrophils, which by itself is critical for the resolution of infection (63), and is supported by our observation of vastly increased accumulation of apoptotic neutrophils in *Hmgb1*^ΔLysM^ livers.

Of several genes, interferon-γ was particularly suppressed in *Hmgb1*^ΔLysM^ compared to *Hmgb1*^f/f^ after 24 hours following infection. It has been suggested that during infection with listeria, interferon-γ production, presumably by NK cells or T lymphocytes, regulates macrophage activation and bacterial clearance as well as the induction of long-term protective cellular immunity (64). Strikingly, *RAGE*^-/-^ mice also display decreased levels of interferon-γ compared to wild-type mice during listeriosis; however, these mice are generally protected from the injurious effects of the microorganism. In line with this notion, RAGE deficiency in dendritic cells, but not in other myeloid cells, resulted in increased bacterial killing compared to *Ager*-floxed controls (**Suppl. Fig. S7**), thus RAGE-induced early inflammation may be deleterious in early stages of infection, yet may ultimately promote protective adaptive Th1 cellular immunity.

Finally, our experiments with mixed bone-marrow chimera suggest a framework in which bone-marrow derived cells (i.e., PMNs), which are recruited to the liver in large numbers early after infection, signal to inflammatory monocytes via HMGB1 to attain their recruitment and induction of transcriptional changes required to mount a sufficient innate immune response. The (attenuated) increased susceptibility of *Hmgb1*^f/f^ >*Hmgb1*^ΔLysM^ to infection, however, also argues for HMGB1 signaling from tissue-resident to circulating immune cells during infection. It conclusively appears that HMGB1 signaling is a dynamic function of time and pathogen burden, with parenchymal cells, tissue-resident and circulating immune cells each signaling to each other via HMGB1 in context-dependent manners.

In summary, we report a vitally important role for HMGB1 in the immunological clearance of the gram-positive bacterium *Listeria monocytogenes*. Our results indicate that while immune cell functions are preserved in the absence of intracellular HMGB1, myeloid cells communicate via HMGB1 during the early phase of infection to direct immune cell migration to the liver, promote their survival and mount signaling pathways that are critical for timely and effective immune responses to the pathogen. Parenchymal cell HMGB1 may actively participate in the immune response only when pathogen burden and tissue damage exceed certain thresholds, and then fuel inflammatory responses. Additional experimental insights into the complex and context-dependent effects of HMGB1 are needed to precisely identify clinical contexts, in which modulation of HMGB1 signaling may confer a benefit for an organism.

## Methods

### Ethics Statement

All animal experiments were assessed and approved by the ethics committee of the Behoerde für Gesundheit und Verbraucherschutz of the City of Hamburg (permit no. 42/15), and conducted under applicable German law governing the care and use of animals for scientific research (Tierschutzgesetz §7 and §8).

### Animals

Generation of *Hmgb1*- and *Ager*-floxed animals was described previously (33,65). *Hmgb1*^f/f^ and *Ager*^f/f^ mice were crossed with albumin-Cre (24), lysozyme-Cre (27) or Cd11c-Cre (66) mice (all from Jackson labs), respectively, to attain cell-specific knockdown of HMGB1 or RAGE. All animals used in this study were on a C57BL/6 background and housed under specific pathogen-free (SPF) conditions in individually ventilated cages with standard food and water ad libitum on a 12-hour day/night cycle. For all experiments, age- and sex-matched control mice from the same colony were used. Mice were infected with 2×10^4^ wildtype *Listeria monocytogenes* strain EGD in 200 μl sterile PBS via the lateral tail vein, and analyzed at the specified time points post-infection (p.i.). In some experiments, mice received daily i.p.-injections of 100 μg anti-HMGB1 or IgG control antibodies, respectively, for three consecutive days of the infection. To determine bacterial titers, 200-300mg of the left hepatic lobe were weighed and mechanically homogenized in 5 ul/g 0.1% Triton X-100 in H_2_0, and suspensions were serially diluted. Dilutions were plated on TSB-agar plates and incubated at 37°C. Colony-forming units (CFU) were counted the next day, and bacterial density in tissues was calculated. Where applicable, samples from the same livers were used for histology, RNA and protein extractions.

### FACS analysis

Cell analysis was performed using a BD LSRFortessa™ cell analyzer (BD Biosciences, USA). For the analysis, single color stainings of each fluorophore (antibodies shown in **Suppl. Table 1**) as well as unstained samples were prepared in order to compensate the fluorescent channels. Cellular events were acquired using BD FACSDiva™ software (BD Biosciences, USA) and subsequently analyzed using FlowJo software (Tree Star, Inc., USA). Gating strategies are shown in **Supplemental Figure S8**.

### Western blot, immunohistochemistry, ELISA and RNA expression analysis

IL1β-, IFNγ-, (R&D Systems) and HMGB1 ELISAs (IBL International) were performed according to the manufacturers’ instructions. RNA from snap-frozen tissues was column-purified using NucleoSpin® RNA kit (Macherey Nagel GmbH und Co. KG, Germany). Following reverse transcription, qPCR was performed using TaqMan primer-probe pairs (Applied Biosystems, USA) and normalized to 18S by comparative Ct (ΔΔC_T_, primers shown in **Suppl. Table 2**). Electrophoresis of protein extracts and subsequent blotting were performed as previously described (5). Blots were incubated with rabbit antibody to HMGB1 (Abcam, ab18256), LC3 (Cell Signaling, 2755) and p62 (Abcam, ab109012) at a dilution of 1:500-1:4000 and visualized by the enhanced chemiluminescence light method (Pierce, USA). Blots were reprobed with mouse antibody to β-actin (Sigma, A5441) or α-Tubulin (Cell Signaling, 3873). Immunohistochemical staining was performed on paraffin-embedded liver sections after 4% formalin fixation using primary antibodies against CD45 (Abcam, ab10558), F4/80 (Biolegend, 123102), HMGB1 (Abcam, ab18256) and *Listeria monocytogenes* (Abcam, ab35132), respectively, followed by HRP-linked anti-rabbit or anti-rat IgG, and developed with DAB Peroxidase substrate (Dako, USA). For immunofluorescence stainings, tissue samples were fixed using 4% PFA followed by 30% sucrose and frozen. Stainings were performed using antibodies against Ly6G (PE-labeled, Biolegend, 127608), p62 (Abcam, ab109012) and *Listeria monocytogenes* (Abcam, ab35132), followed by fluorescently labeled secondary antibodies (Thermo Fisher Scientific, USA). TUNEL staining was performed using the *In Situ* Cell Death Detection Kit (Roche, Germany) according to the manufacturer’s instructions. Microscopy was performed on a Keyence BZ-X710 microscope (Keyence, Japan), with a 10x or 40x lens, respectively.

### Bone marrow transplantation

Bone marrow transplantation (BMT) experiments were performed as previously described (5). Briefly, 4×10^6^ bone marrow cells from donor animals were intravenously injected into lethally irradiated (2×6Gy) recipients. Infection with *Listeria monocytogenes* was performed 4 weeks after bone marrow transplantation.

### Generation and stimulation of bone marrow-derived macrophages (BMDM) and primary polymorphonuclear neutrophils (PMN)

Bone marrow was isolated from femur and tibia of naive mice. For the differentiation of BMDMs, cells were resuspended in growth medium (MEM alpha medium + 10% fetal calf serum, 5% antibiotic/antimycotic (Thermo Fisher Scientific, USA), 10 ng/μl M-CSF (Peprotech, USA), and seeded onto 12-well plates. After seven days of differentiation with M-CSF, bone marrow-derived macrophages were left untreated or primed using IFNγ (0.01 μg/ml) for 16 hours at 37°C before subsequent challenge. BMDMs were washed three times with PBS to remove antibiotics from the cells and subsequently infected with *Listeria monocytogenes* (MOI = 10), and incubated for 1 h at 37°C. Infected cells were then washed three times with PBS and suspended in fully supplemented medium containing Gentamicin to neutralize remaining extracellular bacteria and assess intracellular bacterial degradation. After 30 min, cells were again washed with PBS. Cells in one part of the wells were then lysed using 0.1 % Triton X-100 in ddH_2_O and lysates were plated on TSB agar plates. This time point is considered as 0 h. Remaining wells containing cells were incubated for another 2 and 4 h, respectively, with fully-supplemented medium without antibiotics and then lysed and plated as the first time point. Agar plates were then incubated at 37°C overnight and bacterial colonies subsequently quantified. In seperate experiments, BMDMs were incubated in medium containing lipopolysaccharide (LPS; 1μg/ml) for 4 hours before analysis. For the isolation of neutrophils, red blood cells were lysed using hypotonic saline, then bone marrow cells were separated using a Histopaque gradient (Thermo Fisher Scientific, USA). Cells were seeded onto 96-well plates and rested for 1 hour at 37 °C. Both BMDMs and neutrophils were stimulated with live or heat-inactivated *Listeria monocytogenes* (MOI = 10; neutrophil killing MOI = 0.05), respectively, as indicated in the respective paragraphs and figures. For monocyte phagocytosis assays, BMDMs were primed with 50 ng/ml LPS for 24 hours to attain a M1 phenotype prior to pathogen exposure.

### Nanostring RNA expression and nCounter data analysis

Analysis was performed on liver samples from the indicated groups using the nCounter^®^ SPRINT Profiler (NanoString Technologies) and the nCounter mouse myeloid innate immunity panel V2, which contains 754 unique gene barcodes in 19 pathways across 7 different myeloid cell types. Probes for several housekeeping genes such as ribosomal ribosomal protein L19 (RPL19) are incorporated in the Nanostring codeset and were used for analysis along with positive and negative controls. RNA was prepared and run according to the manufacturer’s protocol. RNA was loaded at 50 ng per sample, and no low-count quality control flags were observed for any of the samples. Data normalization and differential expression analysis was carried out with nSolver analysis software version 4.0 (NanoString Technologies Inc., USA). Genes with an FDR < 0.05 or p-value < 0.05 for the comparison within control groups/injected groups, respectively, were considered being significantly differentially expressed. Visualization of the normalized counts of differentially expressed genes was performed using the statistical framework R (v3.5.1) and an over-representation enrichment analysis for GeneOntology-terms (Biological Process) was carried out with WebGestalt (vfcc27621) (67). Additionally, the significantly differentially expressed genes were visualized as gene interaction networks with Cytoscape (v3.7.1) (68). The underlying interactions of that network were obtained from the STRING database (v11.0) (69). The complete dataset has been deposited at Dryad for reviewing purposes (https://datadryad.org/stash/share/jfT_wz5AB7FrKhPjVcQzLkuwo6CfodmAOM4eehEWx88).

### Statistics

All data are expressed as mean ± standard error of the mean (SEM). For comparison of two groups, Mann-Whitney test was used. For multiple groups, Kruskal-Wallis test with Dunn’s post-test was used. A p-value < 0.05 was considered statistically significant.

## Author contributions

A.V. conducted experiments, analyzed data and provided important intellectual input. K.F., K.L., K.B., S.K. and M.B. conducted experiments, technical assistance. M.Q.: bioinformatic analysis of Nanostring data. M.N., K.R. and R.F.S.: contribution of reagents or mice and intellectual input. A.W.L, S.H. and H.W.M.: intellectual input and revision of the manuscript. P.H.: acquisition of funding, study oversight, data analysis and manuscript preparation.

## Acknowledgements

*Ager*-floxed mice were a kind gift from Prof. Dirk Arnold, DKFZ Heidelberg. This study was supported by German Research Foundation (DFG) grant HU1953/2-1 and the Clinician-Scientist Program of the University Medical Center Hamburg-Eppendorf (to P.H.).

## Supporting Information Legends

**Supplementary Fig S1:** Cre-lox-mediated deletion of HMGB1. (A) *Hmgb1* gene and loxP-sites. (B) Deletion strategy. (C) HMGB1 immunohistochemistry of liver sections from untreated *Hmgb1*^*f/f*^ and *Hmgb1*^Δhep^ mice. (D) HMGB1 protein levels in extracts from whole liver and bone-marrow derived macrophages (BMDM), respectively, from mice of the indicated genotypes. (E). HMGB1 mRNA levels in livers (n=3 per group) and BMDM (n=4 per group). (F) Immunofluorescence for HMGB1 and F4/80 expression in liver sections of untreated *Hmgb1*^*f/f*^ and *Hmgb1*^ΔLysM^ mice. (G) HMGB1 qPCR from isolated Kupffer cells (n=4 isolates per group). BMDM = bone-marrow derived monocytes. (E), (G): Mann-Whitney test. *p<0.05, * * * p<0.001 Bar=100μm.

**Supplementary Fig S2:** Exacerbated disturbances of splenic tissue architecture, accumulation of splenic bacteria and levels of key circulating cytokines Interferon gamma and IL-1β in *Hmgb1*^ΔLysM^ mice. (A) H&E-stained sections of spleens from *Hmgb1*^f/f^ and *Hmgb1*^ΔLysM^ at the indicated time points following infection. (B) Immunohistochemistry for *Listeria monocytogenes* in the spleens of *Hmgb1*^f/f^ and *Hmgb1*^ΔLysM^ mice 72 h after infection. (C) Serum levels of IL-1β and Interferon-γ in *Hmgb1*^f/f^ and *Hmgb1*^ΔLysM^ following infection with 2×10^4^ *Listeria monocytogenes* (n=6-8 per group). Bars=200μm (10x) and 50μm (40x), respectively. h.p.i.=hours post-infection. (C): Mann-Whitney test. n.s.=statistically non-significant. * * p<0.01.

**Supplementary Fig S3:** Accumulation and turnover of F4/80-positive liver macrophages. (A) F4/80 immunohistochemistry of *Hmgb1*^*f/f*^ and *Hmgb1*^ΔLysM^ mice at the indicated time points (n=6-7 per group). (B) Hepatic F4/80 expression assessed by whole liver-qPCR (n=6 per group). (B): Mann-Whitney test. * *p<0.01, Bars = 50μm (10x) and 200μm (40x), respectively.

**Supplementary Fig S4:** Impaired recruitment of hepatic immune cells following infection with less virulent *Listeria monocytogenes*. (A) Hepatic bacterial titers 24 hours and 72 hours after i.v.-injection with a batch of 2×10^4^ *Listeria monocytogenes* (n=6 per group) with attenuated virulence. (B) IFN-γ serum levels in both groups of mice 24h after infection. (C) CD45 immunohistochemistry of whole liver sections and (D) density analysis of hepatic CD45^+^ cells 24 hours after infection. (A)-(D): Mann-Whitney test. n.s.= statistically non-significant. Bars = 200μm.

**Supplementary Fig S5:** HMGB1 deficiency in dendritic cells impairs bacterial clearance from the liver. (A) Hepatic bacterial titers in *Hmgb1*^f/f^ and *Hmgb1*^ΔDC^ mice at the indicated time points (n=5-7 per group). (B) FACS analysis of live intrahepatic immune cells. (C) FACS analysis of live hepatic infiltrating immune cells. (D) H&E- and anti-HMGB1-stainings in liver sections of the indicated mice 72h after infection. (E) qPCR of hepatic inflammatory gene induction 24h and 72h after injection of 2×10^4^ Listeria monocytogenes. (A)-(E): Kruskal-Wallis test with Dunn’s post-test. *p<0.05, * *p<0.01, Bars = 200μm and 50μm, respectively.

**Supplementary Fig S6:** Interaction network of differentially expressed genes. Genes which are upregulated in *Hmgb1*^ΔLysM^ vs *Hmgb1*^f/f^ at baseline (upper diagram) and after exposure to Listeria monocytogenes (lower diagram), respectively, are shown in red whereas downregulated genes are blue. The size of the nodes corresponds to the FDR [or p-value], larger size stands for smaller FDR (or p-value). Outer rings indicate whether a gene is associated with significantly overrepresented GeneOntology-terms.

**Supplementary Fig S7: Dendritic-cell-specific RAGE deficiency reduces virulence of listeriosis in mice**. (A) photomicrographs of primary BMDMs of the indicated genotype via bright-field and GFP channel. RAGE-deficient cells are GFP^+^ (cf. ref. (65)). (B) Hepatic bacterial titers in *Ager*^f/f^ and *Ager*^ΔLysM^ 72 hours after infection (n = 11-14 per group). (C) Hepatic bacterial titers in *Ager*^f/f^ and *Ager*^ΔDC^ 72h after injection of Listeria (n = 5-9 per group). (D) FACS analysis of hepatic cell viability and (E) accumulation of hepatic CD11c^+^ dendritic cells, Cd11b^+^Ly6G^+^ neutrophils and Cd11b^+^Ly6C^+^ monocytes 72 hours after infection. (F) qPCR of hepatic proinflammatory gene expression. (A)-(F) Mann-Whitney test. *p<0.05, * *p<0.01, n.s.=statistically non-significant.

**Supplemental table 1:** FACS antibodies used in the present study

**Supplementary Figure S8:** FACS gating strategies used in the present study.

**Supplemental table 2:** qPCR primers used in the present study

## Notes

**Conflict of interest:** The authors have declared that no conflict of interest exists.

## References

1. Medzhitov R. Origin and physiological roles of inflammation. Nature. 2008 Jul 24;454(7203):428–35.

2. Rock KL, Latz E, Ontiveros F, Kono H. The sterile inflammatory response. Annu Rev Immunol. 2010;28:321–42.

3. Zhang W, Guo S, Li B, Liu L, Ge R, Cao T, et al. Proinflammatory effect of high-mobility group protein B1 on keratinocytes: an autocrine mechanism underlying psoriasis development. J Pathol. 2017 Feb;241(3):392–404.

4. Tsung A, Sahai R, Tanaka H, Nakao A, Fink MP, Lotze MT, et al. The nuclear factor HMGB1 mediates hepatic injury after murine liver ischemia-reperfusion. J Exp Med. 2005 Apr 4;201(7):1135–43.

5. Huebener P, Pradere J-P, Hernandez C, Gwak G-Y, Caviglia JM, Mu X, et al. The HMGB1/RAGE axis triggers neutrophil-mediated injury amplification following necrosis. J Clin Invest. 2015 Feb;125(2):539–50.

6. Yuan H, Jin X, Sun J, Li F, Feng Q, Zhang C, et al. Protective effect of HMGB1 a box on organ injury of acute pancreatitis in mice. Pancreas. 2009 Mar;38(2):143–8.

7. Sawa H, Ueda T, Takeyama Y, Yasuda T, Shinzeki M, Nakajima T, et al. Blockade of high mobility group box-1 protein attenuates experimental severe acute pancreatitis. World J Gastroenterol. 2006 Dec 21;12(47):7666–70.

8. Venereau E, Casalgrandi M, Schiraldi M, Antoine DJ, Cattaneo A, De Marchis F, et al. Mutually exclusive redox forms of HMGB1 promote cell recruitment or proinflammatory cytokine release. J Exp Med. 2012 Aug 27;209(9):1519–28.

9. Germani A, Limana F, Capogrossi MC. Pivotal advances: high-mobility group box 1 protein--a cytokine with a role in cardiac repair. J Leukoc Biol. 2007 Jan;81(1):41–5.

10. Liu K, Mori S, Takahashi HK, Tomono Y, Wake H, Kanke T, et al. Anti-high mobility group box 1 monoclonal antibody ameliorates brain infarction induced by transient ischemia in rats. FASEB J. 2007 Dec;21(14):3904–16.

11. Wang H, Bloom O, Zhang M, Vishnubhakat JM, Ombrellino M, Che J, et al. HMG-1 as a late mediator of endotoxin lethality in mice. Science. 1999 Jul 9;285(5425):248–51.

12. Yang H, Ochani M, Li J, Qiang X, Tanovic M, Harris HE, et al. Reversing established sepsis with antagonists of endogenous high-mobility group box 1. Proc Natl Acad Sci USA. 2004 Jan 6;101(1):296–301.

13. Deng M, Tang Y, Li W, Wang X, Zhang R, Zhang X, et al. The Endotoxin Delivery Protein HMGB1 Mediates Caspase-11-Dependent Lethality in Sepsis. Immunity. 2018 16;49(4):740-753.e7.

14. Andersson U, Tracey KJ. HMGB1 is a therapeutic target for sterile inflammation and infection. Annu Rev Immunol. 2011;29:139–62.

15. Stevens NE, Chapman MJ, Fraser CK, Kuchel TR, Hayball JD, Diener KR. Therapeutic targeting of HMGB1 during experimental sepsis modulates the inflammatory cytokine profile to one associated with improved clinical outcomes. Sci Rep. 2017 19;7(1):5850.

16. Andersson U, Yang H, Harris H. Extracellular HMGB1 as a therapeutic target in inflammatory diseases. Expert Opin Ther Targets. 2018;22(3):263–77.

17. Tang D, Kang R, Livesey KM, Kroemer G, Billiar TR, Van Houten B, et al. High-mobility group box 1 is essential for mitochondrial quality control. Cell Metab. 2011 Jun 8;13(6):701–11.

18. Yanai H, Matsuda A, An J, Koshiba R, Nishio J, Negishi H, et al. Conditional ablation of HMGB1 in mice reveals its protective function against endotoxemia and bacterial infection. Proc Natl Acad Sci USA. 2013 Dec 17;110(51):20699–704.

19. Pamer EG. Immune responses to Listeria monocytogenes. Nat Rev Immunol. 2004 Oct;4(10):812–23.

20. Wang D, Liu K, Wake H, Teshigawara K, Mori S, Nishibori M. Anti-high mobility group box-1 (HMGB1) antibody inhibits hemorrhage-induced brain injury and improved neurological deficits in rats. Sci Rep. 2017 10;7:46243.

21. Fu L, Liu K, Wake H, Teshigawara K, Yoshino T, Takahashi H, et al. Therapeutic effects of anti-HMGB1 monoclonal antibody on pilocarpine-induced status epilepticus in mice. Sci Rep. 2017 26;7(1):1179.

22. Hernandez C, Huebener P, Pradere J-P, Antoine DJ, Friedman RA, Schwabe RF. HMGB1 links chronic liver injury to progenitor responses and hepatocarcinogenesis. J Clin Invest. 2018 Jun 1;128(6):2436–51.

23. Khambu B, Huda N, Chen X, Antoine DJ, Li Y, Dai G, et al. HMGB1 promotes ductular reaction and tumorigenesis in autophagy-deficient livers. J Clin Invest. 2018 Jun 1;128(6):2419–35.

24. Postic C, Shiota M, Niswender KD, Jetton TL, Chen Y, Moates JM, et al. Dual roles for glucokinase in glucose homeostasis as determined by liver and pancreatic beta cell-specific gene knock-outs using Cre recombinase. J Biol Chem. 1999 Jan 1;274(1):305–15.

25. Evankovich J, Cho SW, Zhang R, Cardinal J, Dhupar R, Zhang L, et al. High mobility group box 1 release from hepatocytes during ischemia and reperfusion injury is mediated by decreased histone deacetylase activity. J Biol Chem. 2010 Dec 17;285(51):39888–97.

26. Blériot C, Dupuis T, Jouvion G, Eberl G, Disson O, Lecuit M. Liver-resident macrophage necroptosis orchestrates type 1 microbicidal inflammation and type-2-mediated tissue repair during bacterial infection. Immunity. 2015 Jan 20;42(1):145–58.

27. Clausen BE, Burkhardt C, Reith W, Renkawitz R, Förster I. Conditional gene targeting in macrophages and granulocytes using LysMcre mice. Transgenic Res. 1999 Aug;8(4):265–77.

28. Wang G, Lin A, Han Q, Zhao H, Tian Z, Zhang J. IFN-γ protects from apoptotic neutrophil-mediated tissue injury during acute Listeria monocytogenes infection. Eur J Immunol. 2018;48(9):1470–80.

29. Tang D, Kang R, Livesey KM, Cheh C-W, Farkas A, Loughran P, et al. Endogenous HMGB1 regulates autophagy. J Cell Biol. 2010 Sep 6;190(5):881–92.

30. Huang H, Nace GW, McDonald K-A, Tai S, Klune JR, Rosborough BR, et al. Hepatocyte-specific high-mobility group box 1 deletion worsens the injury in liver ischemia/reperfusion: a role for intracellular high-mobility group box 1 in cellular protection. Hepatology. 2014 May;59(5):1984–97.

31. Kang R, Tang D, Schapiro NE, Loux T, Livesey KM, Billiar TR, et al. The HMGB1/RAGE inflammatory pathway promotes pancreatic tumor growth by regulating mitochondrial bioenergetics. Oncogene. 2014 Jan 30;33(5):567–77.

32. Zhu X, Messer JS, Wang Y, Lin F, Cham CM, Chang J, et al. Cytosolic HMGB1 controls the cellular autophagy/apoptosis checkpoint during inflammation. J Clin Invest. 2015 Mar 2;125(3):1098–110.

33. Huebener P, Gwak G-Y, Pradere J-P, Quinzii CM, Friedman R, Lin C-S, et al. High-mobility group box 1 is dispensable for autophagy, mitochondrial quality control, and organ function in vivo. Cell Metab. 2014 Mar 4;19(3):539–47.

34. Klionsky DJ, Abdelmohsen K, Abe A, Abedin MJ, Abeliovich H, Acevedo Arozena A, et al. Guidelines for the use and interpretation of assays for monitoring autophagy (3rd edition). Autophagy. 2016;12(1):1–222.

35. Homma T, Ishibashi D, Nakagaki T, Satoh K, Sano K, Atarashi R, et al. Increased expression of p62/SQSTM1 in prion diseases and its association with pathogenic prion protein. Sci Rep. 2014 Mar 28;4:4504.

36. Lee H-M, Yuk J-M, Kim K-H, Jang J, Kang G, Park JB, et al. Mycobacterium abscessus activates the NLRP3 inflammasome via Dectin-1-Syk and p62/SQSTM1. Immunol Cell Biol. 2012 Jul;90(6):601–10.

37. Zheng YT, Shahnazari S, Brech A, Lamark T, Johansen T, Brumell JH. The adaptor protein p62/SQSTM1 targets invading bacteria to the autophagy pathway. J Immunol. 2009 Nov 1;183(9):5909–16.

38. Witter AR, Okunnu BM, Berg RE. The Essential Role of Neutrophils during Infection with the Intracellular Bacterial Pathogen Listeria monocytogenes. J Immunol. 2016 01;197(5):1557–65.

39. Serbina NV, Salazar-Mather TP, Biron CA, Kuziel WA, Pamer EG. TNF/iNOS-producing dendritic cells mediate innate immune defense against bacterial infection. Immunity. 2003 Jul;19(1):59–70.

40. Serbina NV, Kuziel W, Flavell R, Akira S, Rollins B, Pamer EG. Sequential MyD88-independent and -dependent activation of innate immune responses to intracellular bacterial infection. Immunity. 2003 Dec;19(6):891–901.

41. Bianchi ME. HMGB1 loves company. J Leukoc Biol. 2009 Sep;86(3):573–6.

42. Liu T, Zhang L, Joo D, Sun S-C. NF-κB signaling in inflammation. Signal Transduction And Targeted Therapy. 2017 Jul 14;2:17023.

43. Cooper MA, Fehniger TA, Ponnappan A, Mehta V, Wewers MD, Caligiuri MA. Interleukin-1beta costimulates interferon-gamma production by human natural killer cells. Eur J Immunol. 2001 Mar;31(3):792–801.

44. Masters SL, Mielke LA, Cornish AL, Sutton CE, O’Donnell J, Cengia LH, et al. Regulation of interleukin-1beta by interferon-gamma is species specific, limited by suppressor of cytokine signalling 1 and influences interleukin-17 production. EMBO Rep. 2010 Aug;11(8):640–6.

45. Higgins SJ, Xing K, Kim H, Kain DC, Wang F, Dhabangi A, et al. Systemic release of high mobility group box 1 (HMGB1) protein is associated with severe and fatal Plasmodium falciparum malaria. Malar J. 2013 Mar 19;12:105.

46. Lee H-W, Kim T-S, Kang Y-J, Kim J-Y, Lee S, Lee W-J, et al. Up-regulated S100 calcium binding protein A8 in Plasmodium-infected patients correlates with CD4(+)CD25(+)Foxp3 regulatory T cell generation. Malar J. 2015 Oct 5;14:385.

47. Cunha C, Carvalho A, Esposito A, Bistoni F, Romani L. DAMP signaling in fungal infections and diseases. Front Immunol. 2012;3:286.

48. Alleva LM, Budd AC, Clark IA. Systemic release of high mobility group box 1 protein during severe murine influenza. J Immunol. 2008 Jul 15;181(2):1454–9.

49. Tsai S-Y, Segovia JA, Chang T-H, Morris IR, Berton MT, Tessier PA, et al. DAMP molecule S100A9 acts as a molecular pattern to enhance inflammation during influenza A virus infection: role of DDX21-TRIF-TLR4-MyD88 pathway. PLoS Pathog. 2014 Jan;10(1):e1003848.

50. Achouiti A, van der Meer AJ, Florquin S, Yang H, Tracey KJ, van’t Veer C, et al. High-mobility group box 1 and the receptor for advanced glycation end products contribute to lung injury during Staphylococcus aureus pneumonia. Crit Care. 2013 Dec 16;17(6):R296.

51. Jonsson N, Nilsen T, Gille-Johnson P, Bell M, Martling C-R, Larsson A, et al. Calprotectin as an early biomarker of bacterial infections in critically ill patients: an exploratory cohort assessment. Crit Care Resusc. 2017 Sep;19(3):205–13.

52. Sims GP, Rowe DC, Rietdijk ST, Herbst R, Coyle AJ. HMGB1 and RAGE in inflammation and cancer. Annu Rev Immunol. 2010;28:367–88.

53. Wang H, Yang H, Czura CJ, Sama AE, Tracey KJ. HMGB1 as a late mediator of lethal systemic inflammation. Am J Respir Crit Care Med. 2001 Nov 15;164(10 Pt 1):1768–73.

54. Calogero S, Grassi F, Aguzzi A, Voigtländer T, Ferrier P, Ferrari S, et al. The lack of chromosomal protein Hmg1 does not disrupt cell growth but causes lethal hypoglycaemia in newborn mice. Nat Genet. 1999 Jul;22(3):276–80.

55. Kurihara T, Warr G, Loy J, Bravo R. Defects in macrophage recruitment and host defense in mice lacking the CCR2 chemokine receptor. J Exp Med. 1997 Nov 17;186(10):1757–62.

56. Edelson BT, Unanue ER. MyD88-dependent but Toll-like receptor 2-independent innate immunity to Listeria: no role for either in macrophage listericidal activity. J Immunol. 2002 Oct 1;169(7):3869–75.

57. Seki E, Tsutsui H, Tsuji NM, Hayashi N, Adachi K, Nakano H, et al. Critical roles of myeloid differentiation factor 88-dependent proinflammatory cytokine release in early phase clearance of Listeria monocytogenes in mice. J Immunol. 2002 Oct 1;169(7):3863–8.

58. Lutterloh EC, Opal SM, Pittman DD, Keith JC, Tan X-Y, Clancy BM, et al. Inhibition of the RAGE products increases survival in experimental models of severe sepsis and systemic infection. Crit Care. 2007;11(6):R122.

59. Janot L, Secher T, Torres D, Maillet I, Pfeilschifter J, Quesniaux VFJ, et al. CD14 works with toll-like receptor 2 to contribute to recognition and control of Listeria monocytogenes infection. J Infect Dis. 2008 Jul 1;198(1):115–24.

60. Yu M, Wang H, Ding A, Golenbock DT, Latz E, Czura CJ, et al. HMGB1 signals through toll-like receptor (TLR) 4 and TLR2. Shock. 2006 Aug;26(2):174–9.

61. Aucott H, Sowinska A, Harris HE, Lundback P. Ligation of free HMGB1 to TLR2 in the absence of ligand is negatively regulated by the C-terminal tail domain. Mol Med. 2018 04;24(1):19.

62. Tadie J-M, Bae H-B, Jiang S, Park DW, Bell CP, Yang H, et al. HMGB1 promotes neutrophil extracellular trap formation through interactions with Toll-like receptor 4. Am J Physiol Lung Cell Mol Physiol. 2013 Mar 1;304(5):L342–349.

63. Poon IKH, Lucas CD, Rossi AG, Ravichandran KS. Apoptotic cell clearance: basic biology and therapeutic potential. Nat Rev Immunol. 2014 Mar;14(3):166–80.

64. Harty JT, Bevan MJ. Specific immunity to Listeria monocytogenes in the absence of IFN gamma. Immunity. 1995 Jul;3(1):109–17.

65. Constien R, Forde A, Liliensiek B, Gröne H-J, Nawroth P, Hämmerling G, et al. Characterization of a novel EGFP reporter mouse to monitor Cre recombination as demonstrated by a Tie2 Cre mouse line: Characterization of EGFP Reporter Mouse Line. Genesis. 2001 May;30(1):36–44.

66. Caton ML, Smith-Raska MR, Reizis B. Notch-RBP-J signaling controls the homeostasis of CD8-dendritic cells in the spleen. J Exp Med. 2007 Jul 9;204(7):1653–64.

67. Wang J, Vasaikar S, Shi Z, Greer M, Zhang B. WebGestalt 2017: a more comprehensive, powerful, flexible and interactive gene set enrichment analysis toolkit. Nucleic Acids Res. 2017 Jul 3;45(W1):W130–7.

68. Shannon P, Markiel A, Ozier O, Baliga NS, Wang JT, Ramage D, et al. Cytoscape: a software environment for integrated models of biomolecular interaction networks. Genome Res. 2003 Nov;13(11):2498–504.

69. Szklarczyk D, Franceschini A, Wyder S, Forslund K, Heller D, Huerta-Cepas J, et al. STRING v10: protein-protein interaction networks, integrated over the tree of life. Nucleic Acids Res. 2015 Jan;43(Database issue):D447–452.

